# A novel CAR T cell blend targeting PDPN and GD2 to overcome glioblastoma heterogeneity

**DOI:** 10.1101/2025.02.03.636223

**Authors:** Vera Nickl, Julian Hübner, Marah Alsalkini, Rhonda McFleder, Julia Wagner, Renato Liguori, Björn Grams, Moritz Meyer-Hofmann, Leopold Diener, Camelia Maria Monoranu, Carsten Hagemann, Fulvia Ferrazzi, Chi Wang Ip, Ralf-Ingo Ernestus, Hermann Einsele, Michael Hudecek, Mario Löhr, Thomas Nerreter

## Abstract

**Background:** While chimeric antigen receptor (CAR) T cells have achieved encouraging remission rates in hematological malignancies, they demonstrated limited success in treating Glioblastoma (GBM), particularly due to high intra- and intertumoral heterogeneity. In this study, we identified a relevant and preserved target antigen, Podoplanin (PDPN), and evaluated the potential of a PDPN- and GD2-CAR T cell blend to overcome GBM heterogeneity.

**Methods:** Target antigen screening included clinical samples, healthy tissues and cell lines, as well as publicly available RNA sequencing datasets. The anti-tumor function of CAR T cells were examined in co-culture experiments with GBM cell lines and patient-derived organoids (PDOs), and *in vivo* after locoregional delivery in orthotopic xenograft models.

**Results:** The generated CAR T cells demonstrated strong anti-tumor activity against several cell lines and PDOs from multiple patients. PDPN and GD2 expression was detectable in all PDOs at varying densities and regardless of the antigenic profile, the CAR T cell blend induced significantly higher levels of apoptosis in organoids than single antigen targeting counterparts. *In vivo*, we observed efficient tumor regression after locoregional administration of monospecific CAR T cells. While heterogeneous orthotopic tumors eventually relapsed in these groups, blended therapy resulted in a significantly increased overall survival and even achieved cure in the majority of mice.

**Conclusion:** This novel PDPN-/GD2-CAR T cell blend demonstrated strong efficacy in advanced preclinical models of glioblastoma. The results suggest that this approach can overcome GBM heterogeneity in clinical application and address previous limitations of single antigen CAR T cell therapies.

**KEYPOINTS:**

- PDPN is a relevant and consistent CAR T cell target antigen in primary and recurrent GBM
- PDPN- and GD2-CAR T cells display synergistic anti-tumor activity in patient-derived organoids
- Local delivery of our CAR T cell blend confers long-term survival and cure in GBM-bearing mice

**IMPORTANCE OF THE STUDY:** The highly variable landscape of tumor antigens in GBM represents a serious obstacle to single antigen CAR T cell therapies. In this study, we identified Podoplanin (PDPN) and GD2 as the most preserved target antigens in a screening campaign, with an even increased density in recurrent GBM compared to primary GBM. We combined PDPN- and GD2-CAR T cells in a blended treatment approach to counter antigen heterogeneity, and achieved a substantial gain in efficacy against patient-derived organoids (PDOs). In an orthotopic xenograft model, locoregional administration of the PDPN- and GD2-CAR T cell blend conferred long-term complete remission and increased survival compared to single antigen targeting. This study provides a hierarchy of CAR target antigens for treating GBM and illustrates the therapeutic potential of combinatorial antigen targeting using a PDPN/GD2-CAR T cell blend.

**GRAPHICAL ABSTRACT:**
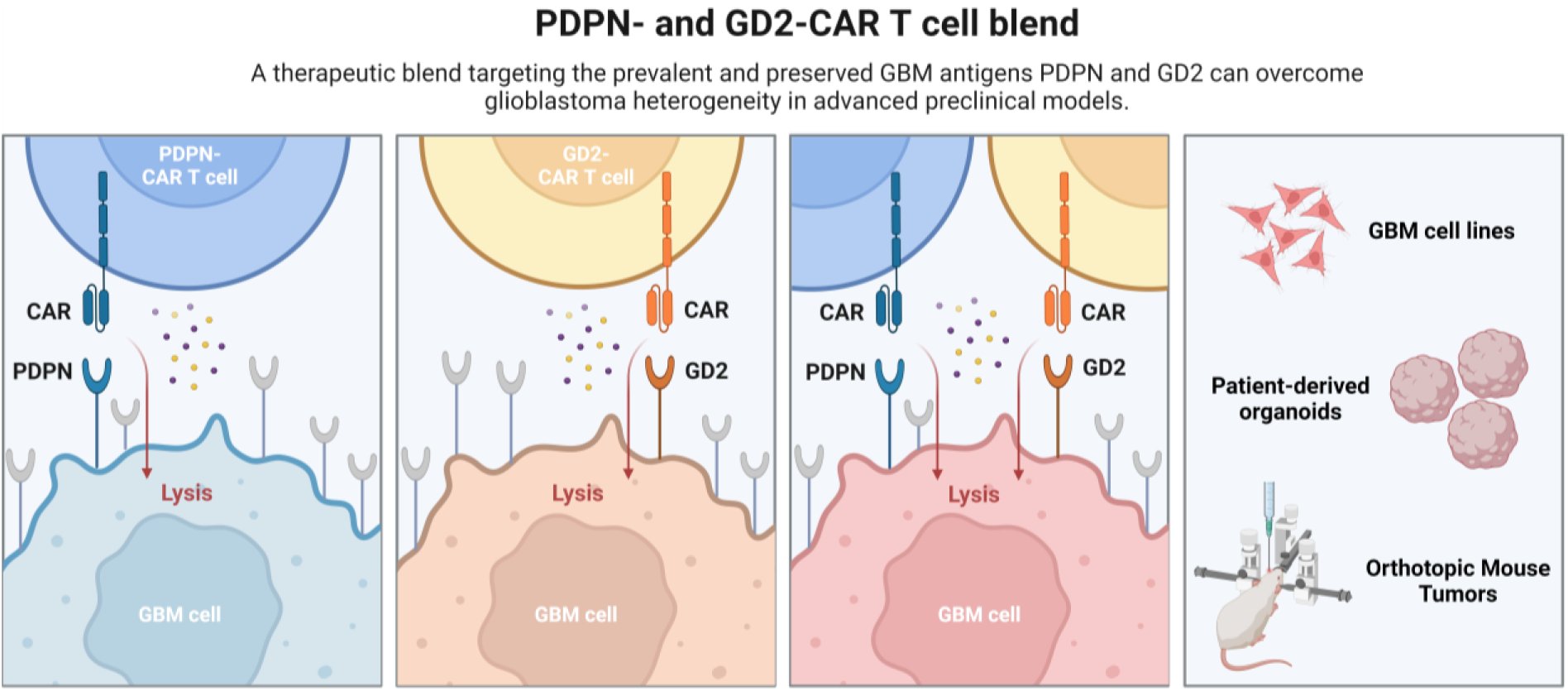

## INTRODUCTION

Glioblastoma (GBM) is the most common and most aggressive primary malignant brain tumor in adults and is considered incurable despite extensive treatment.^1,2^ Median survival after standard therapy, consisting of surgical resection followed by radiotherapy with concomitant and adjuvant chemotherapy with temozolomide (TMZ) and tumor treating fields^3^, remains limited to 21 months, with a 5-year survival prognosis of only 13 %.^4–6^

Chimeric antigen receptor (CAR) T cells targeting CD19 have remarkable clinical efficacy in a variety of hematologic neoplasms.^7^ However, effective CAR T cell therapy has not yet been demonstrated in solid tumors, and particularly GBM.^8,9^ The immunosuppressive tumor microenvironment (TME), CAR T cell trafficking to tumor sites and especially the high degree of inter- and intratumoral heterogeneity present major challenges to CAR T cell therapy.^10,11^ Accordingly, phase I clinical trials demonstrated partial responses in recurrent GBM following therapy with e.g. EGFRvIII-, IL13Rα2-, and HER2-CAR T cells, but failed to induce durable remissions.^12–14^

The objectives of this study were to identify and validate antigens that are consistent among patients, and to establish a dual targeting CAR T cell approach capable of overcoming the challenge of heterogeneity. For this purpose, we characterized antigen densities in GBM and identified Podoplanin (PDPN), a mucin-like type-I integral membrane glycoprotein involved in tumorigenesis and metastasis, as lead candidate. Elevated expression levels of PDPN have been found in a number of cancers, particularly those that originate from immune-privileged organs like testis and brain. This overexpression is associated with tumor malignancy, treatment resistance and a dismal prognosis.^15–17^ The second most promising candidate emerging from our screening campaign was GD2, a tumor-specific disialoganglioside that is highly expressed on tumors of neuroectodermal origin and has been validated as a potent and tolerable target antigen for CAR T cell therapy.^18–20^

We evaluated the efficacy of PDPN- and GD2-CAR T cells in experiments with multiple GBM cell lines, patient-derived organoids (PDOs) and an orthotopic GBM model. Generated directly from intraoperatively obtained tumor tissue, PDOs preserve the complexity and heterogeneity of the paternal tumor and represent a model closer to the patient than monolayer cultures.^21^ On the other hand, orthotopic xenograft models including locoregional delivery of CAR T cells into the lateral ventricle represent a physiologically suitable environment for the assessment of CAR T functionality. In these preclinical GBM models, dual targeting with a PDPN- and GD2- CAR T cell blend demonstrated superior antitumor function and achieved durable tumor regression and cure in the majority of mice.

## METHODS

### Glioblastoma tissue

Local ethic committee approval was obtained (#22/20-me) and the study was conducted according to the declaration of Helsinki. Fresh GBM tissue for FACS analysis and PDO generation was obtained intraoperatively after patients provided informed consent to the study. Paraffin-embedded sections were provided by the Department of Neuropathology, University of Würzburg.

### Cell lines

Glioblastoma cell lines (GCLs) DKMG, U373, and U251 were purchased from the German Collection of authenticated Cell Cultures (DSMZ, Leibniz, Germany). Cell lines U138, U343 and U87 were purchased from the European Collection of authenticated Cell Cultures (ECACC, Porton Down, England). GCLs were cultured in Dulbecco’s Modified Eagle’s Medium (DMEM) containing 1 g/L glucose, sodium pyruvate, 3.7 g/L NaHCO_3_ and 4 mM L-glutamine (Gibco, Carlsbad, CA, USA), supplemented with 10 % v/v heat-inactivated fetal calf serum (FCS), 2× nonessential amino acids (NEAA, 100× stock), 1% penicillin/streptomycin (Invitrogen, Carlsbad, CA, USA) as a monolayer in 75 cm^3^ flasks (Corning) at 37 °C, 5 % CO_2_ and 95 % humidity.

### Generation of patient derived organoids

Organoids were prepared as previously described^22^: Briefly, acute tumor tissue was temporarily stored on ice in Hibernate A medium (Gibco). The tumor tissue was cleared from necrosis and blood vessels, then carefully cut into pieces with an McIlwain Tissue Chopper (Cavey Laboratory Engineering, Guildford, UK), treated with RBC Lysis Buffer (Invitrogen) for 10 min and washed two times with Hibernate A medium containing 1 % Glutamax, 0.4 % penicillin/streptomycin and 0.1 % Amphotericin (all from Gibco). Sections were transferred into GBO medium consisting of a 1:1 mixture of DMEM/F12 and Neurobasal media, supplemented with 2 % B27 without Vitamin A (50×), 1 % Glutamax, 1 % NEAA, 0.4 % penicillin/streptomycin, 0.1 % β-Mercaptoethanol (all from Gibco) and 0.023 % human insulin (Sigma-Aldrich, Merck KGaA, Darmstadt, Germany) to ultra-low attachment 6-well plates (Corning, New York, NY, USA) and incubated at 37 °C, 5 % CO_2_ and 95 % humidity on an orbital shaker at 120 rpm. After 2 weeks of culture, organoids adapted a spherical shape, and could be used for experimental procedures.

### Preparation of CAR T cells

CAR T cells were generated as described earlier.^23^ Lentiviral transduction was performed with vectors encoding second generation PDPN- or GD2-CARs comprising a single-chain fragment variable derived from NZ-1 or hu14.18, respectively, an IgG4 spacer and a signaling module of 4-1BB_CD3ζ.

### Orthotopic xenograft model

All animal experiments were approved by the local authorities (Würzburg, Germany). Female six- to eight-week old NSG mice (NOD.Cg-Prkdc^scid^ Il2rg^tm1Wjl^/SzJ (JAX Stock number 005557)) were stereotactically injected on day 0 with 2×10^5^ U87 ffluc/GFP^+^ (firefly luciferase/green fluorescent protein^+^) GBM cells into the caudate nucleus of the right hemisphere at 1.8 mm lateral, 0.5 mm anterior to the bregma and 3.5 mm below the dura. At a rate of 0.1 mm/sec, the needle was advanced 3.8 mm before injection, paused there for 1 min to allow the tissue to bounce back and then moved back 0.3 mm to create a pocket for the tumor cells. A total of 5 µL of cell suspension was injected at a rate of 1 µL/min before the needle was retracted with 1 mm/min in order to avoid efflux of the suspension. Tumor engraftment was verified under anesthesia with isoflurane at day 6 via bioluminescence imaging on an IVIS Lumina (PerkinElmer, Waltham, MA, USA) 10 min after intraperitoneal (i.p.) injection of D-luciferin (Biosynth, Staad, Schweiz) at 0.3 mg per g body weight. Imaging data were analyzed using LivingImage software (PerkinElmer). On day 7, 2×10^6^ CAR T cells or untransduced T cells (1:1 mixture of CD4^+^ and CD8^+^ T cells) were injected i.c.v. into the lateral ventricle of the left hemisphere at 0.95 mm medial, 0.22 mm posterior and 2.3 mm below the dura. A total of 5 µl of cell suspension was injected at a rate of 1 µl/min before the needle was retracted with 1 mm/min. Tumor reduction was subsequently analyzed via bioluminescence imaging and overall survival of the mice was monitored.

### Immunohistochemistry

For IHC, tissue was cut to 2 µm thick slices, deparaffinized in xylene and hydrated in a graded series of alcohols. Heat induced retrieval was either performed for 20-40 min with citrate buffer pH = 6.0 (EGFRvIII, GD2, HER2, PDPN) or for 5 min with Tris-EDTA buffer pH = 9.0 (CD70, CD133, CSPG4, EphA2, IL13Rα2). After blocking with 0.7 % hydrogen peroxide (Sigma-Aldrich), slides were treated with 10 % normal goat serum (Invitrogen) and the primary antibody diluted with antibody buffer (DCS Innovative Diagnostik Systeme, Hamburg, Germany) was applied overnight at 4 °C (see Supplementary data). Slides were then incubated with the secondary antibody for 30 min and labeled with streptavidin peroxidase (Super Sensitive Link-Label IHC Detection System, DCS/BioGenex, Hamburg, Germany). Diaminobenzadine (Liquid DAB+ Substrate Chromogen System, Dako, Wiesentheid, Germany) was applied for 5 min and after rinsing with water, cell nuclei were counterstained using hemalum solution acid (Carl Roth GmBH, Karlsruhe, Germany) before slides were mounted.

Five representative areas of view per slide were photographed with a Leica DMI3000 B light microscope, LEICA DFC450 camera and LAS V4.5 software (all Leica Microsystems, Wetzlar, Germany) with standardized settings at 40x amplification and analyzed for staining intensity via the batch processing function of the open source program Fiji.^24^ The percentage of low, medium and high antigen expressing cells on each slide were quantified by applying a custom macro. The expression score was further weighted by multiplication with 1 (low), 2 (medium) or 3 (high expression). Finally, median expression values of all tumors combined for each antigen were calculated. All analysis was monitored and double-checked with an experienced neuropathologist.

### Flow cytometry

15 patients were included at the Department of Neurosurgery, University Hospital Würzburg. Cell surface expression of target antigens was analyzed by FACS. Acute tissue was cleaned from vessels, surrounding brain tissue as well as debris, homogenized mechanically, washed, resuspended in 100 µL PBS/0.5 % bovine serum albumin (Thermo Fisher Scientific) and after Fc-receptor blocking reagent (Human TruStain FcX™, Biolegend, San Diego, CA, USA) treatment for 5 min stained with 0.5 µL of the respective antibody for 20 min at 4 °C (see Supplementary data). If necessary, secondary staining was performed after washing for 20 min at 4 °C.

Mouse brains were harvested directly after euthanasia with CO_2_, cut into small pieces and subsequently digested using the Tumor Dissociation Kit, human (Miltenyi) and a gentleMACS Octo Dissociator with Heaters (Miltenyi, Program 37C_h_TDK1). Cell debris was removed from viable cells using Debris Removal Solution (Miltenyi) and red blood cell lysis was performed using BD PharmLyse Buffer (BD). After live/dead staining with Zombie Aqua™ (1:1000, Biolegend) and Fc-receptor blocking reagent treatment, cells were stained with anti-GD2-BV421 (1:250, BioLegend, clone: 14G2A) and anti-PDPN-APC/Cy7 (1:250). Analysis was performed on a FACSCanto II (BD). Data were analyzed using FlowJo software v10.7.1 (BD).

### qPCR

Pooled cDNA of healthy organ (Human MTC™ Panel I & II, Takara Bio Inc., Kusatsu, Japan) and healthy brain tissue (TissueScan, Human Brain cDNA Array, HBRT501, OriGene, Rockyville, MD, USA) was analyzed for PDPN expression. Each sample was analyzed with the StepOnePlus Real-time PCR System using TaqMan^TM^ PDPN Gene Expression Assays (Thermo Fisher Scientific, FAM-MGB_PL, Hs00366766_m1) with GAPDH as endogenous control (Thermo Fisher Scientific, VIC-MGB_PL, Hs99999905_m1). Three technical replicates were performed according to the manufacturer’s instructions.

To rank the obtained healthy tissue expression, PDPN expression was analyzed in 29 GBM patients. Total mRNA was extracted from frozen tissue using the PureLink^TM^ RNA Mini Kit (Thermo Fisher Scientific) and subsequently reverse-transcribed using the High Capacity RNA to cDNA^TM^ Kit (Thermo Fisher Scientific). cDNA concentrations were adjusted to 5 ng/µL with ultra-pure water for storage at −80 °C. qPCRs were performed as described above and the mean ΔCt value of the patients was used as calibrator for healthy tissue expression according to the ΔΔCt-method.

### Survival analysis based on TCGA GBM dataset

Survival analyses relied on a publicly available cohort of human GBMs made available by The Cancer Genome Atlas (TCGA) (https://tcga-data.nci.nih.gov/tcga/). In particular, clinical annotations and gene expression data of the TCGA GBM dataset were downloaded relying on the TCGAbiolinks package (v.2.28.4, https://bioconductor.org/packages/TCGAbiolinks) within R version 4.3.1, applying the following custom filters: experimental.strategy = “RNA-Seq”, workflow.type = “STAR - Counts”, and sample.type = “Primary Tumor”. Raw gene counts were then normalized and transformed using the variance-stabilizing transformation (VST) method implemented in the Bioconductor package DESeq2 (v.1.40.0).^25^ The obtained VST counts were used as expression values in the following survival analyses.

Ganglioside synthase genes *ST8SIA1* and *B4GALNT1* were used as GD2-phenotype predicting signatures^26^ and EGFRvIII had to be excluded from this analysis due to the lack of an underlying variant-specific gene signature. To standardize time measurements, the time to death or last follow-up was converted into months for both deceased and living patients. For each target gene, patients were stratified into high- and low-expression groups relying on the surv_cutpoint function within the R package ‘survminer’ (v. 0.4.9, https://CRAN.R-project.org/package=survminer). For each gene, the difference in survival rates between the two groups was assessed using the log-rank test. In addition, the hazard ratio (HR) and its associated significance p-value were computed using the R package ‘survival’ (v. 3.5.7 https://CRAN.R-project.org/package=survival).

### Cytotoxicity assays

GBM cell lines stably transduced with a ffluc/GFP fusion gene were incubated in triplicate wells at 1×10^4^ cells/well. T cells were added to the respective wells at an effector to target (E:T) ratio of 10:1. D-luciferin substrate (Biosynth) was added to the co-culture to a final concentration of 0.15 mg/mL, and the decrease in luminescence signal in wells that contained target cells and T cells compared to target cells only was measured using a luminometer (Tecan, Crailsheim, Germany).

In PDOs, all apoptotic tumor cells were identified after 20 hours of incubation with CAR T cells at an E:T ratio of 1:4. We embedded triplicates in paraffin and subsequently performed co-staining and analysis as described above with primary antibodies against CC3 (1:400, Cell Signaling, Leiden, Netherlands, clone: Asp175:400), GD2 (1:50, Santa Cruz, clone: 53831) or PDPN (ready-to-use, Zytomed, clone: D2-40). After secondary staining with AlexaFluor488 and AlexaFluor555 (both Invitrogen), slices were mounted with Fluoroshield mounting medium containing DAPI (Abcam) and analyzed as described above.

### Cytokine secretion

5×10^4^ T cells were plated in triplicate wells with target cells at an E:T of 4:1, and secretion of IFNγ was measured by ELISA (ELISA MAX, Biolegend) in supernatant removed after 24 hours of incubation. Analysis was performed using a luminometer (Tecan).

### Proliferation assays

T cells were labeled with 0.2 μM carboxyfluorescein succinimidyl ester (CFSE, Invitrogen), washed, and plated in triplicate wells with irradiated (80 Gy) target cells at an E:T ratio of 4:1. No exogenous cytokines were added to the culture medium. After 72 hours of incubation, cells were analyzed by flow cytometry to assess cell division of T cells.

### Statistical analyses

Analyses were performed using GraphPad Prism 9 (GraphPad Software, San Diego, CA, USA). Data was examined for Gaussian distribution by Kolmogorov-Smirnov testing before significance testing was conducted. Experiments with cell lines, primary tissue or organoids were either analyzed by paired t-tests or Two-Way ANOVAs with Sidak’s multiple comparison tests. *In vivo* data was analyzed by One-Way-ANOVA followed by Dunn’s multiple comparison test and Log-rank (Mantel-Cox) test for the Kaplan-Meier survival analysis.

## RESULTS

### PDPN is a relevant and preserved target in GBM

We screened the antigen densities of potential targets in GBM using freshly resected intraoperative tissue from patients with confirmed GBM diagnosis (n=15). Flow cytometric analyses revealed that only PDPN was preserved at high densities in every sample, followed by GD2 with a higher variance (**Figure 1A**).

**Figure 1:**
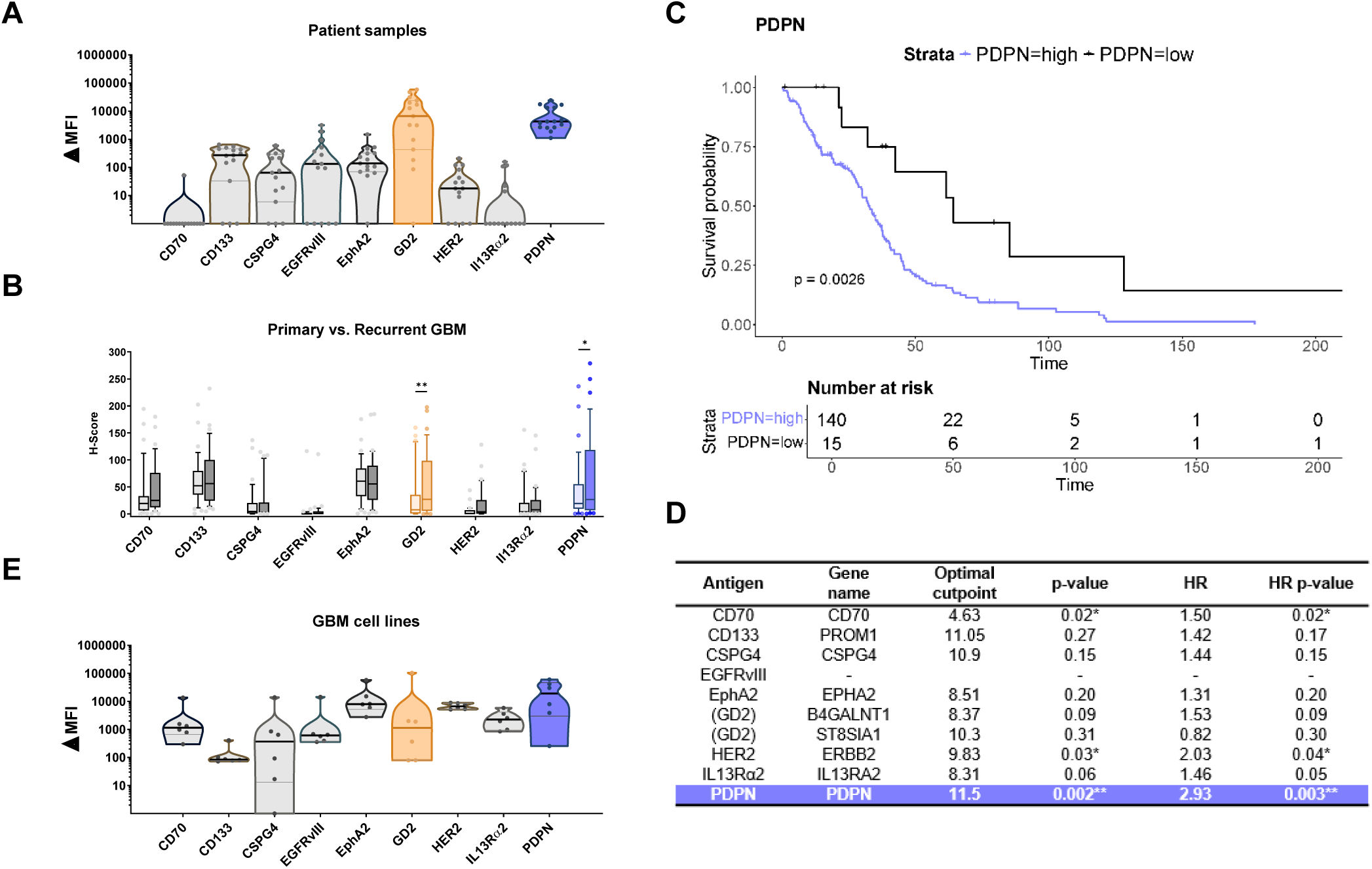
Antigen screening experiments. **(A)** ΔMFIs of primary tissue samples (n = 15). Dark bars represent the median, light bars represent the quartiles. **(B)** Antigen expression in primary (light) and recurrent (dark) GBM specimens (n=35). Median and quartiles are shown. Whiskers represent the 10^th^ – 90^th^ percentile and the points represent statistical outliers. Significant changes in antigen expression (GD2 and PDPN) are marked with an asterisk (*, p<0.05; **, p<0.01). **(C)** Kaplan-Meier survival curve of TCGA-GBM patient cohorts stratified on the basis of PDPN expression (p = log-rank test p-value). **(D)** Results of the analysis of the association between overall survival and the expression of selected genes (Optimal cutpoint: expression cutpoint achieving the strongest difference between survival curves; p-value: log-rank test p-value; HR: hazard ratio; HR p-value: p-value corresponding to the HR). EGFRvIII had to be excluded from this analysis due to the lack of an underlying variant-specific gene signature. **(E)** ΔMFIs of GBM antigens are displayed logarithmically as violin plots for (n = 6) GCLs (DKMG, U87, U138, U251, U343, U373). Dark bars represent the median, light bars represent the quartiles.

To assess the presence of target antigens during disease progression, we analyzed tumor specimens of primary GBM (pGBM) and corresponding recurrent GBM (rGBM) of 35 relapsed patients via immunohistochemistry (IHC) staining. In contrast to all other antigens, exhibiting no difference between the primary tumor and its recurrence, GD2 and PDPN densities were significantly increased at relapse, suggesting both markers as potential candidates for treating recurrent GBM (**Figure 1B**).

To evaluate the impact of the presence of GBM antigens on the overall survival (OS) of patients, we relied on the GBM dataset from the Cancer Genome Atlas (TCGA) and found a clear association between higher PDPN expression and lower OS (**Figure 1C**). Importantly, PDPN expression showed the most significant inverse correlation with patient survival compared to other antigens (**Figure 1D**).

To serve as a predictive model system, GBM cell lines (GCLs) should ideally reflect antigen densities present on primary tissue. Therefore, we performed flow cytometric analyses of all markers on a panel of GCLs (DKMG, U87, U138, U251, U343, U373) and detected each antigen at varying densities, with PDPN showing the highest overall median expression (**Figure 1E**). Interestingly, several antigens preserved on GCLs (e.g. CD70, IL13Rα2) were not consistently expressed on primary tissue as well, in contrast to GD2 and PDPN (**Figure 1A, E**).

To assess potential off-target expression, we performed qPCRs on pooled cDNA from healthy organs. Ovarian, placental, and lung tissues were highest among organs, but overall PDPN expression was low compared to GBM (**Supplementary Figure S1A**). Although expression in pan-brain cDNA was negligible, we performed additional qPCR analysis with pooled cDNA of 24 distinct healthy brain regions to further assess the generally low expression of PDPN in healthy tissue (**Supplementary Figure S1B**). While some brain areas exhibited minor expression of PDPN on cDNA level, IHC analysis showed no signal of PDPN in healthy brain tissue on protein level. Additionally, sections from different tumor locations revealed mutable expression patterns with low, medium and high levels of all tested antigens, reflecting the high intratumoral heterogeneity of GBM (**Supplementary Figure S1C**).

These findings demonstrate that PDPN is a relevant and consistent target antigen in GBM with an expression pattern predestined for CAR T cell therapy.

### Synergy of PDPN- and GD2-specific CAR T cells *in vitro*

In light of our findings, we equipped T cells with a second-generation CAR comprising a single-chain fragment variable (scFv) derived from the PDPN-specific monoclonal antibody (mAb) NZ-1 (**Figure 2A**). Purification, gene transfer rates and overall expansion were highly efficient for both CD4^+^ and CD8^+^ CAR T cells (**Figure 2B-D**), yielding a well-balanced T cell product consisting mainly of central memory-like (T_CM_), effector memory-like (T_EM_), and stem cell memory-like (T_SCM_) phenotypes (**Figure 2E-G**). Co-culture of PDPN-CAR T cells with GCLs led to efficient and specific target cell lysis in a 24-hour bioluminescence-based cytotoxicity assay (**Figure 2H–J**). Similarly, surface expression of PDPN stimulated proliferation and significantly enhanced secretion of IFNγ in response to the PDPN^+^ cell lines (**Figure 2K, L**).

**Figure 2:**
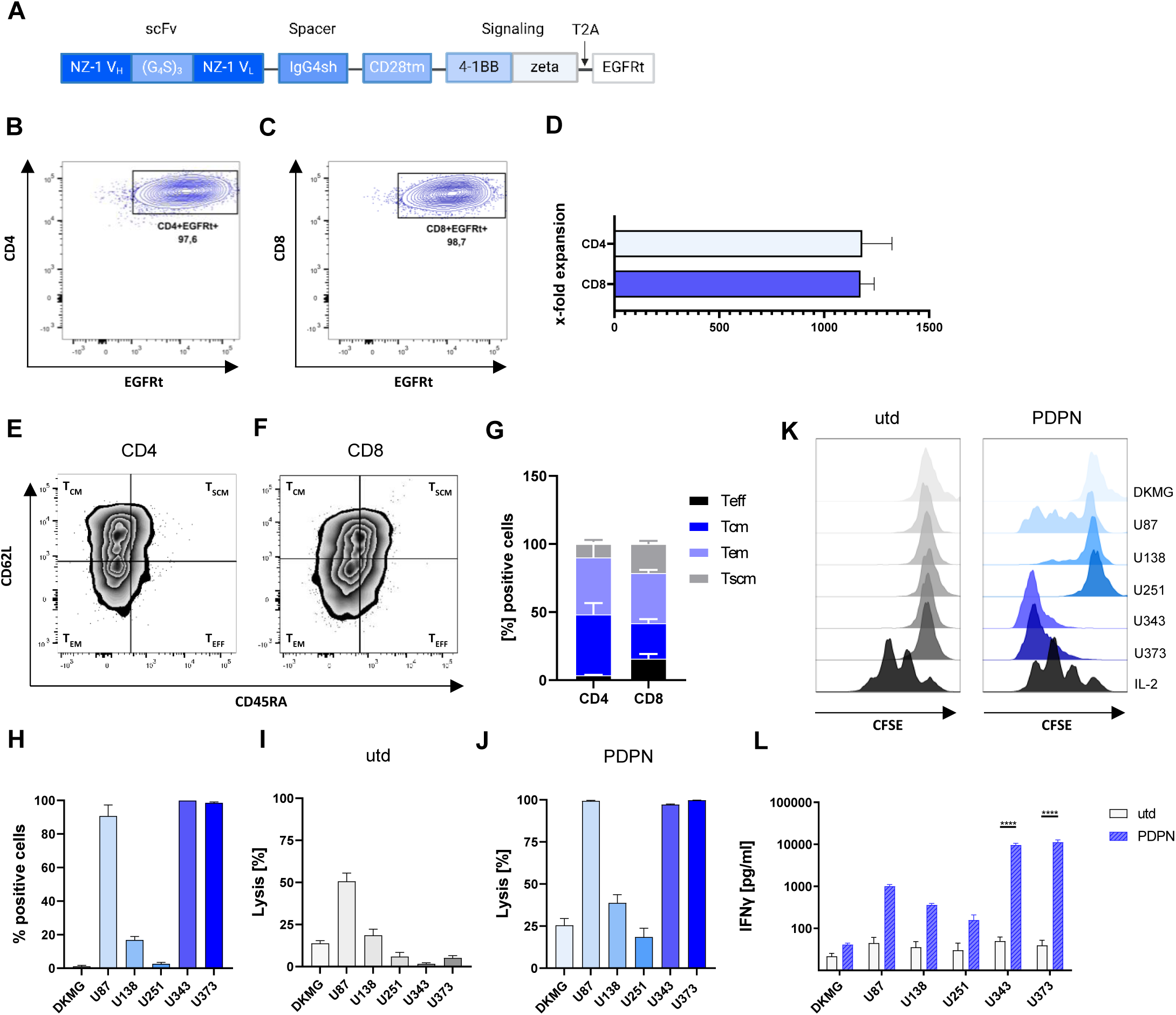
PDPN-CAR T cells are functional against tumor cell lines *in vitro*. **(A)** Structural features of the PDPN-specific CAR construct. Created with BioRender.com. **(B, C)** Representative purity of CD4^+^ and CD8^+^ CAR T cells. **(D)** Enrichment of CAR^+^ T cells in a 10- day expansion protocol. **(E-G)** Representative memory-like phenotypes of expanded CD4^+^ and CD8^+^ CAR T cells. **(H)** Percentage of PDPN-positive cells in GCLs (DKMG, U87, U138, U251, U343, U373) determined by flow cytometry. **(I, J)** Tumor cell lysis after incubation with untransduced T cells and PDPN-CAR T cells after 24 hours. **(K)** A representative CFSE assay shows the proliferation of CD8^+^ CAR T cells after 72 hours of co-culture with GCLs. 250 U/µl IL-2 was used as a positive control. **(L)** ELISA-based quantification of IFNγ from supernatants collected after 24 hours of incubation with GCLs. Data are shown as mean + SEM (****, p<0.0001).

The expression pattern of GD2 suggested its suitability as combinatory antigen in a blended CAR T cell approach (**Figure 1A-C**). Therefore, we generated CAR T cells based on a scFv fragment derived from the GD2-specific antibody 14.18. Purification, gene transfer rates and T cell expansion were equally efficient for GD2-and PDPN-CAR T cells (**Figure 2B-D** and **Supplementary Figure S2A-D**). Notably, antigen staining, cytotoxicity and cytokine analysis indicated synergistic functionality of GD2- and PDPN-CAR T cells. Particularly, in 24-hour assays, five of the six GCLs were efficiently lysed by GD2- or PDPN-CAR T cells alongside significant IFNγ production (**Supplementary Figure S2E-H and Figure 2J, L**).

### CAR T cell blend consistently demonstrates superior efficacy in PDOs

To investigate potential synergy in a heterogeneous model representing primary tumor characteristics, we conducted dual targeting experiments with PDOs. CAR T cells generated from healthy donors were co-incubated with PDOs derived from tissue of eight consecutive patients at an inverse effector-to-target (E:T) ratio of 1:4 (**Figure 3A**), whereas PDPN- and GD2-CAR T cells were mixed in a 1:1 ratio for blended therapy (**Figure 3B**). In co-culture experiments with CAR T cells, complete degradation occurred after 48 hours of incubation, complicating evaluation using immunofluorescence microscopy (IF). Therefore, we selected an interim analysis time point of 20 hours for organoid harvest and processing (**Figure 3C, D**).

**Figure 3:**
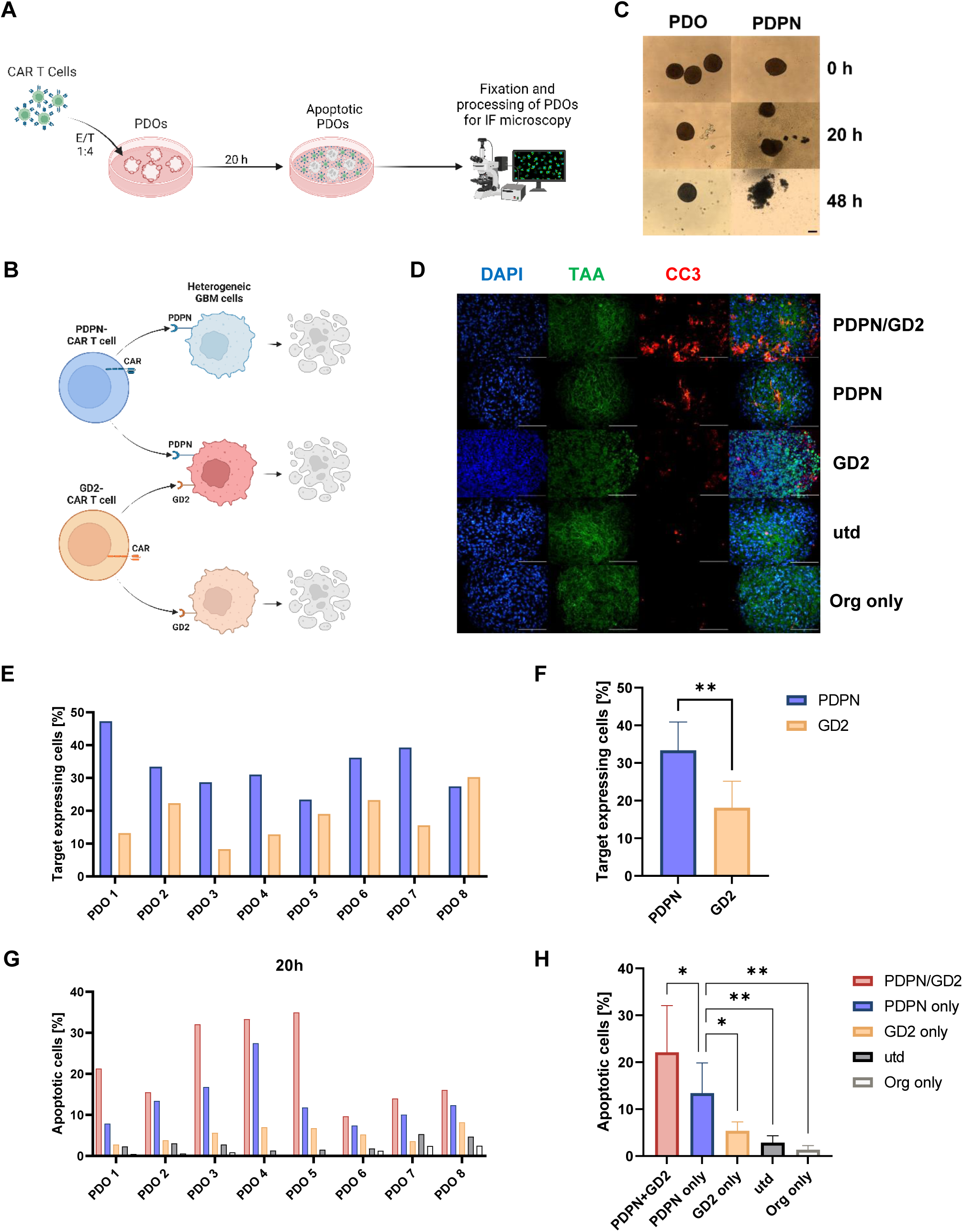
Dual targeting PDPN/GD2 CAR T cells induce significantly higher levels of apoptosis in PDOs. **(A)** CAR T cells were co-cultured with PDOs at an inverse effector-to-target ratio of 1:4 for 20 hours and PDOs were fixed and processed for IF microscopy. Created with BioRender.com. **(B)** The independently generated PDPN- and GD2-CAR T cells were mixed in a 1:1 ratio for dual targeting therapy of heterogeneous tumor cells. **(C)** Representative light microscopy pictures of organoids without (PDO) and with CAR T cell treatment (PDPN) at different time points. Bright field mode with standard settings was used. Scale bar: 50 μm. **(D)** Representative IF images of tumor cell apoptosis (DAPI+/TAA+/CC3+, TAA = tumor-associated antigen). **(E, F)** IF-based analysis of PDPN and GD2 expression in PDOs generated from eight consecutive donors. **(G, H)** IF-based analysis of tumor cell apoptosis after 20 hours of co-incubation. All experiments were performed in triplicates. Data are shown as mean + SEM (*, p<0.5; **, p<0.01). Magnification: 40x. Scale bar: 50 µm.

Both antigens were detectable in all PDOs at varying densities, demonstrating conserved but heterogeneous expression of both markers across primary tissue of different patients (**Figure 3E, F**). PDPN-CAR T cells induced significantly higher levels of apoptosis compared to GD2-CAR T cells, consistent with a higher level of expression of PDPN in the PDOs. Importantly, dual treatment proved to be the best therapeutic modality in each individual set of PDOs, regardless of antigen densities (**Figure 3G**). Combined in a therapeutic blend, both cell products acted synergistically and induced significantly higher levels of tumor cell apoptosis compared to single antigen targeting approaches (**Figure 3D, H**).

Co-culture experiments were repeated with PDOs and T cells generated from the same patients in an autologous setting (**Supplementary Figure S3A**). Peripheral blood mononuclear cells (PBMCs) were isolated following surgery and histopathological confirmation of GBM diagnosis, prior to the start of the patient’s radiochemotherapeutic treatment. PDOs were harvested after 20 hours of co-culture with autologous or allogeneic PDPN- and GD2- CAR T cells and processed for IF analysis. As visualized by representative IF images, CAR T cells induced robust levels of apoptosis with no statistically significant differences between the efficacy of autologous PDPN- or GD2-CAR T cells and those derived from healthy donors (**Supplementary Figure S3B, C**). Furthermore, these results demonstrate the potential of this organoid model for patient-specific preclinical assessment of CAR T cell efficacy prior to therapy.

### Curative dual targeting in an orthotopic xenograft model

To complete our preclinical experiments, we established an orthotopic xenograft model and implanted tumors stereotactically in NSG mice. U87 ffluc^+^/GFP^+^ cells were injected into the caudate nucleus of the right hemisphere on day 0, and tumor engraftment was verified by bioluminescence imaging (BLI) on day 6 (**Figure 4A**). We selected U87 for our xenografts due to its heterogeneous antigen expression pattern in flow cytometric analysis, with populations expressing either GD2, PDPN, both or neither of the markers (**Figure 4B**). Interestingly, U87 tumors isolated from the brains of untreated mice showed variations in the expression pattern, with either an increased GD2 expression or higher levels of PDPN, both or none, suggesting a certain degree of plasticity *in vivo* (**Figure 4C, D**). Therefore, the heterogeneity of these tumors presented a challenging environment for our dual targeting approach.

**Figure 4:**
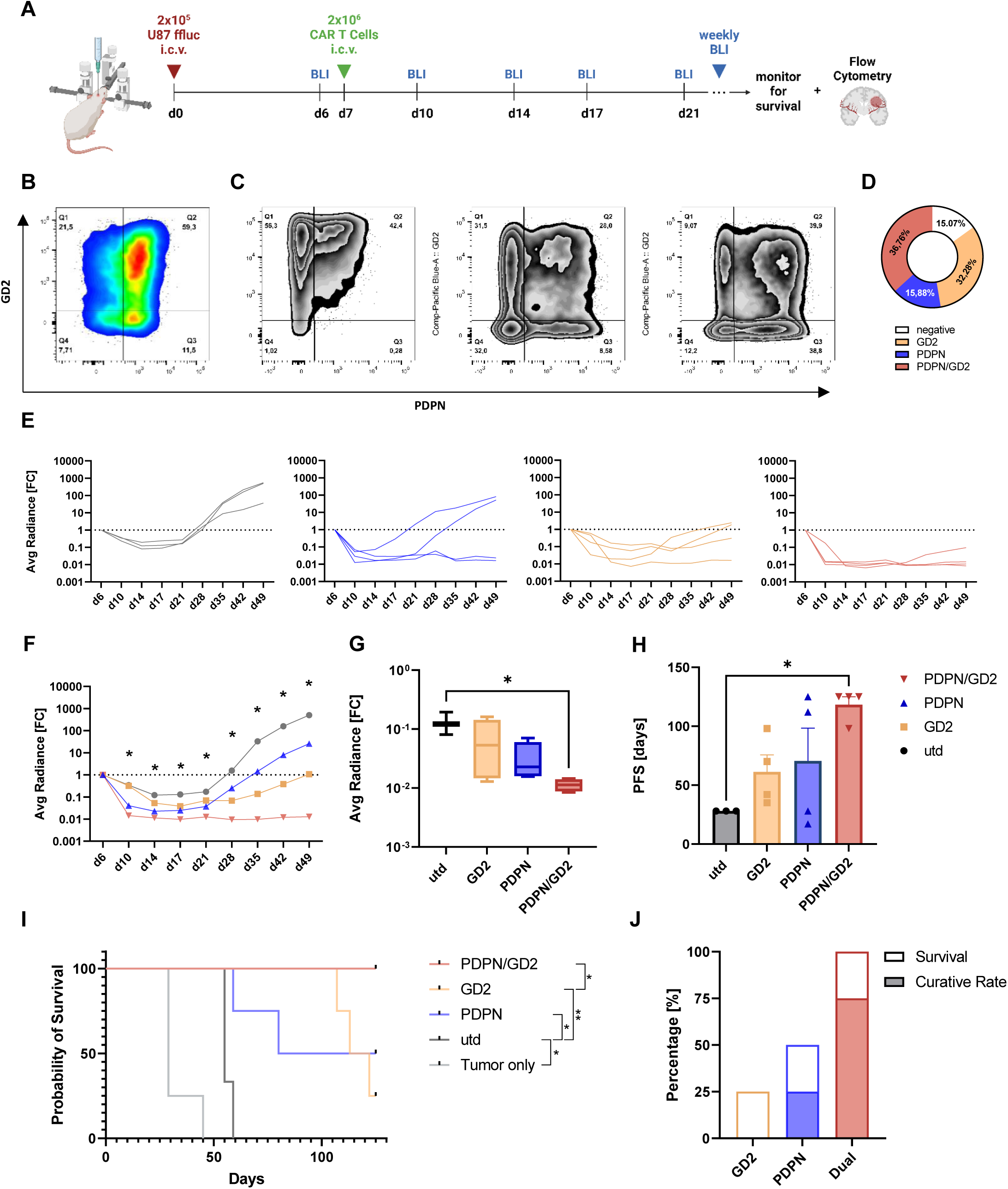
Synergistic effects of a PDPN- and GD2-CAR T cell blend in orthotopic mouse models. **(A)** Female 6- to 8-week old NSG mice were stereotactically injected with 2×10^5^ U87 ffluc/GFP^+^ cells into the caudate nucleus of the right hemisphere on day 0. After verifying the tumor engraftment on day 6 using bioluminescence imaging (BLI), 2×10^6^ CAR T cells or untransduced T cells (1:1 mixture of CD4^+^ and CD8^+^ T cells) were injected intracerebroventricularly (i.c.v.) into the lateral ventricle of the left hemisphere. Tumor reduction and overall survival of the mice were then monitored. Created with BioRender.com. **(B)** Representative PDPN/GD2 expression pattern of U87 cells after standard *in vitro* cultivation analyzed by flow cytometry. **(C-D)** Analysis of the antigen expression patterns of untreated U87 tumors. **(E, F)** Monitoring of the fold change in average radiance over time. Data are shown as median and individual curves of mice. An asterisk indicates time points of significantly reduced tumor load between the dual targeting and utd control groups (*, p<0.05). **(G)** Analysis of initial response depth at day 7 after T cell administration. Median and quartiles are shown. Whiskers represent the 10^th^ – 90^th^ percentile. **(H)** Analysis of progression-free survival (PFS), defined in our model as the time interval of tumor recurrence beyond the initial BLI signal on day 6. Data are shown as mean + SEM (*, p<0.05). **(I)** Kaplan-Meier survival analysis of untreated, untransduced T cell and CAR T cell treated mice (*, p<0.5; **, p<0.01). **(J)** Percentage of surviving mice and curative rates after treatment with GD2-, PDPN-, or PDPN/GD2-CAR T cells by the end of a 125-day observation period.

As the intrathecal injection of CAR T cells via an Ommaya reservoir appears to be superior to intravenous application^27–29^, we administered our CAR T cells on day 7 directly into the brain, thereby bypassing the blood brain barrier. Injection in the lateral ventricle rather than into the tumor cavity added an additional layer of complexity to CAR T cell trafficking. The reduction in tumor BLI signal and overall survival of mice was subsequently monitored (**Figure 4A**).

Stereotactic injections were successful, evidenced by consistent tumor engraftment in all experimental groups (**Supplementary Figure S4A**). Six weeks after treatment, all mice in the control group met termination criteria. CAR T cell-treated mice exhibited tumor regression and remission following intracerebroventricular (i.c.v.) delivery. However, tumor relapses occurred in mice treated with single antigen CAR T cells, while responses were durable only in mice that received dual treatment with the PDPN- and GD2-targeting blend at the same total CAR T cell dose (**Figure 4E, F**). Assessment of the initial depth of response after T cell administration on day 14 showed that established tumors were already reduced to a certain degree in PDPN- and GD2-CAR T cell groups, but only blended therapy led to a homogeneous and significant reduction (**Figure 4G**). Consistent with this, progression-free survival (PFS), defined in our model as the time interval of tumor recurrence beyond the initial BLI signal on day 6, was significantly increased only in dual treated mice (**Figure 4H**).

With a median survival time of 55 days, already an injection of untransduced T cells provided a survival advantage compared to tumor only mice, consistent with *in vitro* cytotoxicity data (**Figure 2K**). While therapy with either PDPN- or GD2-CAR T cells significantly increased median survival to 103 and 118 days, respectively, the combination of both products in a dual targeting regimen resulted in 100% survival of the treated mice (**Figure 4I and Supplementary Figure S4A, B**). Flow cytometric analysis of the brains of surviving mice confirmed that no GFP^+^ tumor cells could be detected in mice with durable remission **(Supplementary Figure S4C**). Thus, single antigen targeting therapy only led to the cure of one mouse (PDPN group) while in contrast, the CAR T cell blend proved to act synergistically and provided an encouraging curative effect in the majority of the mice (**Figure 4J**).

Collectively, these data demonstrate that the reduction of heterogeneous tumors in a physiological environment following i.c.v. delivery is highly efficient with a dual PDPN/GD2- CAR T cell therapeutic blend.

## DISCUSSION

Given the success of CAR T cells in several hematological malignancies, there have been high expectations for the treatment of solid tumors including GBM. However, single antigen CAR T cell therapies have encountered significant challenges in GBM, including a highly heterogeneous tumor antigen landscape.

In this study, we aimed to target multiple antigens, ideally involved in tumor progression and maintenance, to counteract tumor heterogeneity. For this purpose, we analyzed cell lines, paraffin-embedded GBM specimens and primary tissue using IHC, qPCR, and flow cytometric analyses and investigated the association between expression of the respective antigens and patient survival via a publicly available dataset. Interestingly, we found that established GBM target antigens, e.g. EGFRvIII and IL13Rα2^12,14^, were expressed in all GCLs but only at low levels in our primary tissue sample cohort. In contrast, the type I integral membrane glycoprotein PDPN was expressed at high levels in all tissue samples, and survival analysis showed a clear association between its expression and poor prognosis. PDPN is a marker for tumor lymphatic vessels, and its expression increases with higher glioma WHO grades.^30^ Involved in tumorigenesis, its overexpression promotes phosphorylation of ERM proteins and RhoA activation, which leads to elevated motility and induces epithelial-to-mesenchymal transition.^31,32^ An aggressive subset of glioma stem cells with higher treatment resistance is characterized by PDPN expression and depletion renders glioma cell lines radiosensitive.^33^ Furthermore, glioma cells secrete PDPN in exosomes, leading to PDPN surface expression on macrophages and M2 polarization within the TME after phagocytosis.^34^ Together with the surface expression on cancer-associated fibroblasts within the TME^35^, these findings make PDPN-CAR T cells highly attractive for GBM treatment as they could target both tumor cells and TME components simultaneously.

Previous studies with NZ-1-derived PDPN-CAR T cells did not aim to address GBM heterogeneity or did not result in the cure of mice.^36,37^ Here, we combined CAR T cells targeting PDPN and GD2, a tumor-specific disialoganglioside that is highly expressed in a variety of cancers, but particularly in central nervous system (CNS) tumors.^38–40^ Mainly absent in normal tissues, GD2 is linked to invasion, motility, and proliferation and plays a role in oncogenesis through several signaling pathways.^18,41,42^ GD2 expression was consistently high in GCLs and fresh GBM tissue and besides PDPN, it was the only other antigen among the examined candidates to show significantly increased expression levels in relapsed glioblastoma. Additionally, GD2 is an established CAR T cell antigen and the subject of multiple clinical trials demonstrating feasibility and safety.^20,43^ Given this favorable profile, we chose a dual targeting approach applying a blend of PDPN- and GD2-CAR T cells.

Multiple dual targeting approaches are being investigated, including bicistronic or bispecific CAR designs, Boolean logic gating strategies and simultaneous administration of two separate CAR T cell products.^44–46^ It is still unclear which approach is best suited for which malignancy and antigen combination, as each has advantages and disadvantages, and CAR design and intrinsic signaling contribute to the complexity of multi targeting approaches. To combine both CARs and preserve their encouraging *in vitro* and *ex vivo* functionality in our models, we selected a 1:1 blend of PDPN- and GD2-CAR T cells for this study. In addition, this approach maintains modularity that allows easy addition or combination of other CARs without increasing the complexity of engineering or reducing gene transfer rates caused by a higher genetic cargo.

Cell lines only inadequately represent paternal tumor characteristics due to monolayer cultivation, disruption of cell-to-cell contacts during passaging and the total duration of *in vitro* cultivation. In contrast, PDOs, generated directly from intraoperative primary material within two to four weeks, can partially retain tumor vasculature, glioma stem cell markers, and hypoxic gradients. They represent the primary tumor at histological, molecular, and transcriptional level, even after weeks of culture, freezing, and thawing.^21^ Therefore, PDOs display a heterogeneous and challenging model to study the efficacy of our CAR T cell blend. PDPN and GD2 were detectable at varying levels but in each patient investigated, underlining the conserved yet heterogeneous expression of both antigens. We chose to process organoids for IF analysis within 20 hours of co-culture with CAR T cells to preserve tissue integrity and prevent degradation and loss of organoids after longer incubation times. At this intermediate time point, PDOs treated with untransduced T cells as well as organoid only controls did not disintegrate and retained their spherical shape. In contrast, treatment with single antigen targeting CAR T cells led to profound anti-tumoral effects. However, dual targeting with pooled PDPN- and GD2-CAR T cells was found to be superior in all cases, resulting in significantly increased apoptosis levels in organoids, hereby confirming synergistic functionality.

To evaluate our cell product *in vivo*, we used an advanced orthotopic NSG mouse model, which allows direct assessment of the functionality of our human CAR T cells against human glioblastoma in a physiologically appropriate environment. For this model, we inoculated U87 cells, which showed a high degree of plasticity and heterogeneity *in vivo* and formed different GD2- and PDPN-expressing subpopulations in distinct mice. Despite the lack of a complex tumor microenvironment in NSG mice, our model allowed us to investigate the benefits of dual targeting in reducing heterogeneous tumors following locoregional administration. Rather than systemic or intratumoral delivery, we chose to inject the cell products into the lateral ventricle of mice as clinical studies suggest that direct administration into the cerebrospinal fluid (CSF) is associated with improved treatment efficacy.^47–49^ In a clinical setting, the placement of an Ommaya reservoir in the ventricle allows for repeated administration as well as pharmacodynamic monitoring and rapid control of potentially elevated intracranial pressure and toxicities.^43^ Here, we studied the effect of a single injection on day 7 due to the complexity of the procedures and to minimize surgical burden in the murine model. Nevertheless, locoregional administration of single antigen targeting CAR T cells already resulted in remarkable tumor reduction and partial remission, a result not observed in previous studies applying similar models.^36,50^ Importantly, dual targeting therapy resulted in overall survival and achieved curative effects and durable remissions in the majority of mice. Only one mouse treated with the CAR T blend ultimately relapsed, but survived with a comparatively low tumor burden without symptoms until the end of a 125-day observation period. Given that a single injection elicited such long-lasting responses against heterogeneous U87 tumors, we were able to demonstrate a synergistic and curative effect of pooled PDPN- and GD2-CAR T cells *in vivo*.

Together, we present a novel dual targeting approach with pooled PDPN- and GD2-CAR T cells that efficiently targets various GBM cell lines, heterogeneous patient-derived organoids, and orthotopic xenografts after locoregional delivery. In a clinical application, this novel combination could be used to address GBM heterogeneity and thereby overcome previous limitations of single antigen CAR T cell therapies.

## ACKNOWLEDGEMENT

This work was funded by the Interdisciplinary Center for Clinical Research (IZKF) at the Medical Faculty of the University of Würzburg (Project B450).

Further funding was provided by the Deutsche Forschungsgemeinschaft (DFG, German Research Foundation) –SFB- TRR 338/1 2021 –452881907 (projects A02 to MH and A05 to TN), SFB- TRR 221/2 –324392634 (project A03 to HE&MH) and SFB- TRR 305 –429280966 (project Z01 to FF).

We are grateful to the animal staff at the Center for Experimental Molecular Medicine (ZEMM; University Hospital Würzburg & University of Würzburg) for providing excellent animal care. We further thank Dagmar Hemmerich (Department of Neurosurgery, Section Experimental Neurosurgery, University Hospital Würzburg) for great technical assistance.

## AUTHOR CONTRIBUTIONS

Conceptualization: TN, VN, MH, ML; Methodology and Validation: JH, TN, VN, MH, ML, RMF, CWI, MA, CMM; Software: JH, RL, FF, TN, VN; Formal Analysis: JH, TN, VN, MA; Investigation: JH, MA, JW, BG, VN, LD, MMH; Resources: MH, HE, RIE, CWI, CH; Writing – Original Draft: JH, TN, VN; Writing – Review & Editing: JH, TN, MH, VN, CH; Supervision: TN, VN, MH, ML; Funding Acquisition: TN, VN, MH.

## DECLARATION OF DATA AVAILABILITY

The raw data that support the findings of this study are available from the corresponding authors upon reasonable request.

## SUPPLEMENTARY MATERIAL

### Antibody List

| **Target** | **Clone** | **Company** | **Dilution** |
| --- | --- | --- | --- |
| CD70 | NP_001243.1 | Abcam | 1:800 |
| CD133 | AC133 | Miltenyi | 1:50 |
| CSPG4 | NG2 | Proteintech Group, Inc. | 1:500 |
| EphA2 | SP169 | ZytoMed Systems GmbH | 1:200 |
| EGFRvIII | EGFRvIII.1 | ZytoMed Systems GmbH | 1:100 |
| GD2 | 2Q549 | USBiological | 1:100 |
| Her2 | SP3 | ZytoMed Systems GmbH | ready-to-use |
| IL13Rα2 | Cys22-Leu342 | R&D Systems | 1:200 |
| PDPN | D2-40 | ZytoMed Systems GmbH | ready-to-use |

**FACS**
| **Target** | **Fluorochrome** | **Clone** | **Company** | **Comments** |
| --- | --- | --- | --- | --- |
| CD70 | APC-Vio770 | REA292 | Miltenyi |  |
| CD133 | BV421 | 315-2C11 | Biolegend |  |
| CSPG4 | VioBlue | EP-1 | Miltenyi |  |
| EphA2 | PE | SHM16 | Biolegend |  |
| EGFRvIII |  | DH8.3 | Novus | primary |
| anti-mouse IgG1 | FITC | RMG1-1 | Biolegend | secondary |
| GD2 | BV421 | 14G2A | Biolegend |  |
| GD2 | PE-Cy7 | 14G2A | Biolegend |  |
| Her2 | FITC | 24D2 | Biolegend |  |
| IL13Rα2 | PE-Vio770 | REA308 | Miltenyi | |
| PDPN | APC-Cy7 | NC-08 | Biolegend |  |

**Supplementary Figure S1:**
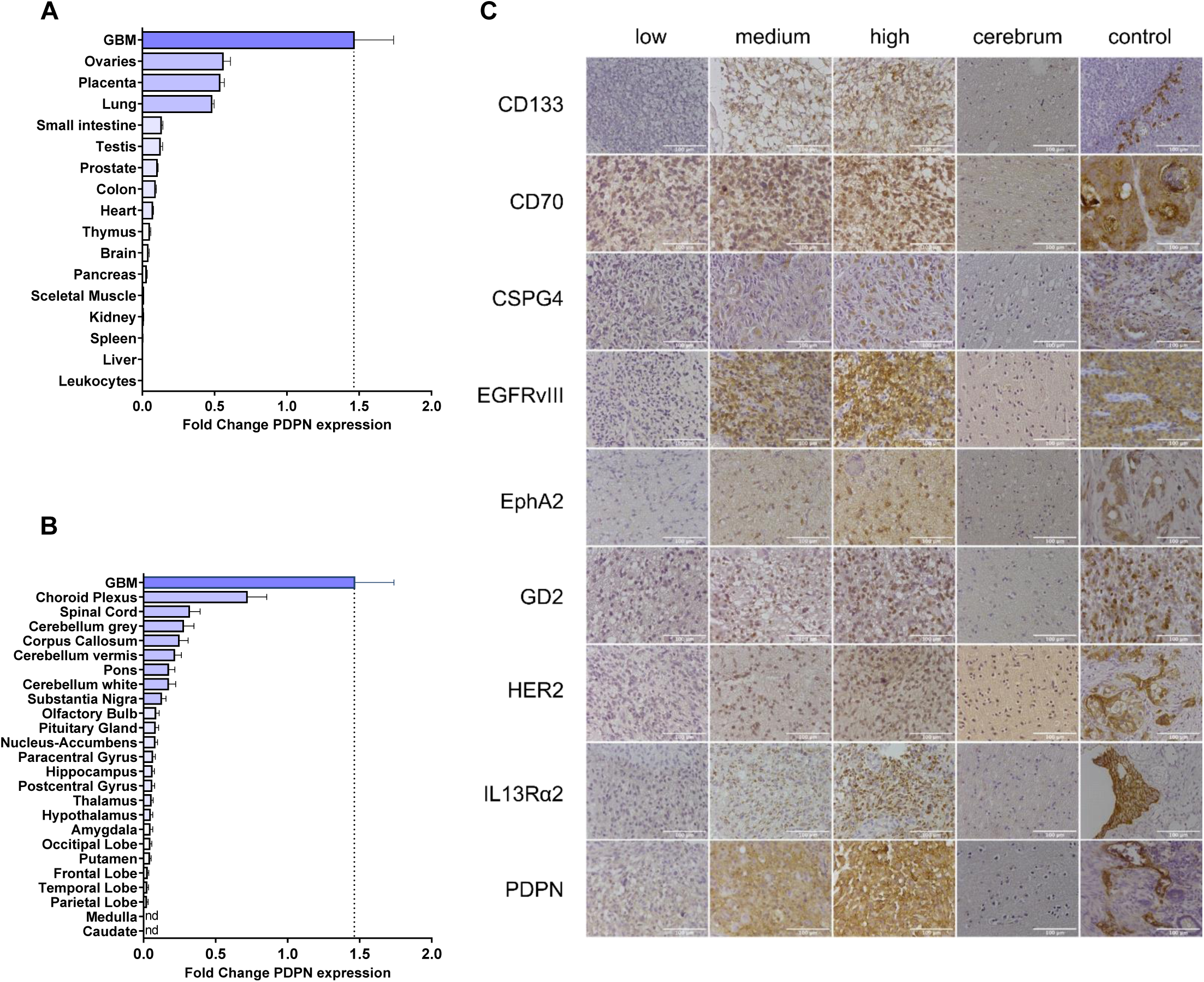
qPCR and IHC analysis of PDPN expression. **(A-B)** Fold change PDPN expression in healthy organ and healthy brain tissue. Mean + SEM are shown. **(C)** For each antigen, areas of low, medium, and high staining intensity were found in the same GBM sample. A healthy brain control (cerebrum) showed no staining; a representative sample of n = 3 healthy donors is shown. Colon carcinoma (CD133), tonsil tissue (CD70), lung carcinoma (CSPG4), EGFRvIII-positive GBM (EGFRvIII), breast carcinoma (EphA2), neuroblastoma (GD2), HER2-positive breast carcinoma (HER2), prostate carcinoma (IL13Rα2), and colon lymphatics (PDPN) were used as positive controls. Magnification: 40x. Scale bar: 100 µm.

**Supplementary Figure S2:**
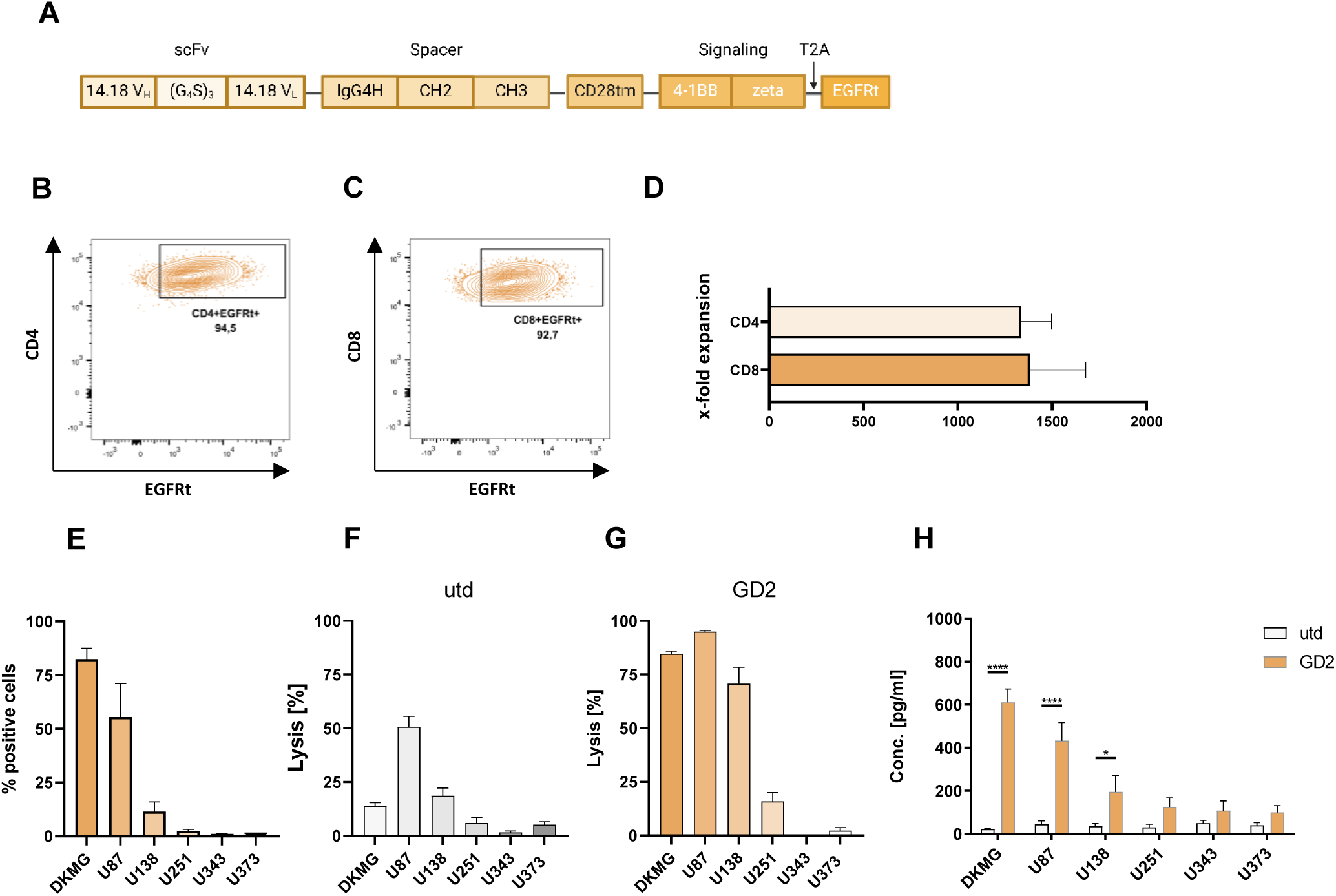
GD2-CAR T cells are functional against tumor cell lines *in vitro*. **(A)** Structural features of the GD2-specific CAR construct. Created with BioRender.com. **(B, C)** Representative purity of CD4^+^ and CD8^+^ CAR T cells. **(D)** Enrichment of CAR^+^ T cells in a 10-day expansion protocol. **(E)** Percentage of GD2-positive cells in GCLs (DKMG, U87, U138, U251, U343, U373) determined by flow cytometry. **(F, G)** Tumor cell lysis after incubation with utd or GD2-CAR T cells after 24 hours. **(H)** ELISA-based quantification of IFNγ from supernatants collected after 24 hours of incubation with GCLs. Data are shown as mean + SEM (*, p<0.05; ****, p<0.0001).

**Supplementary Figure S3:**
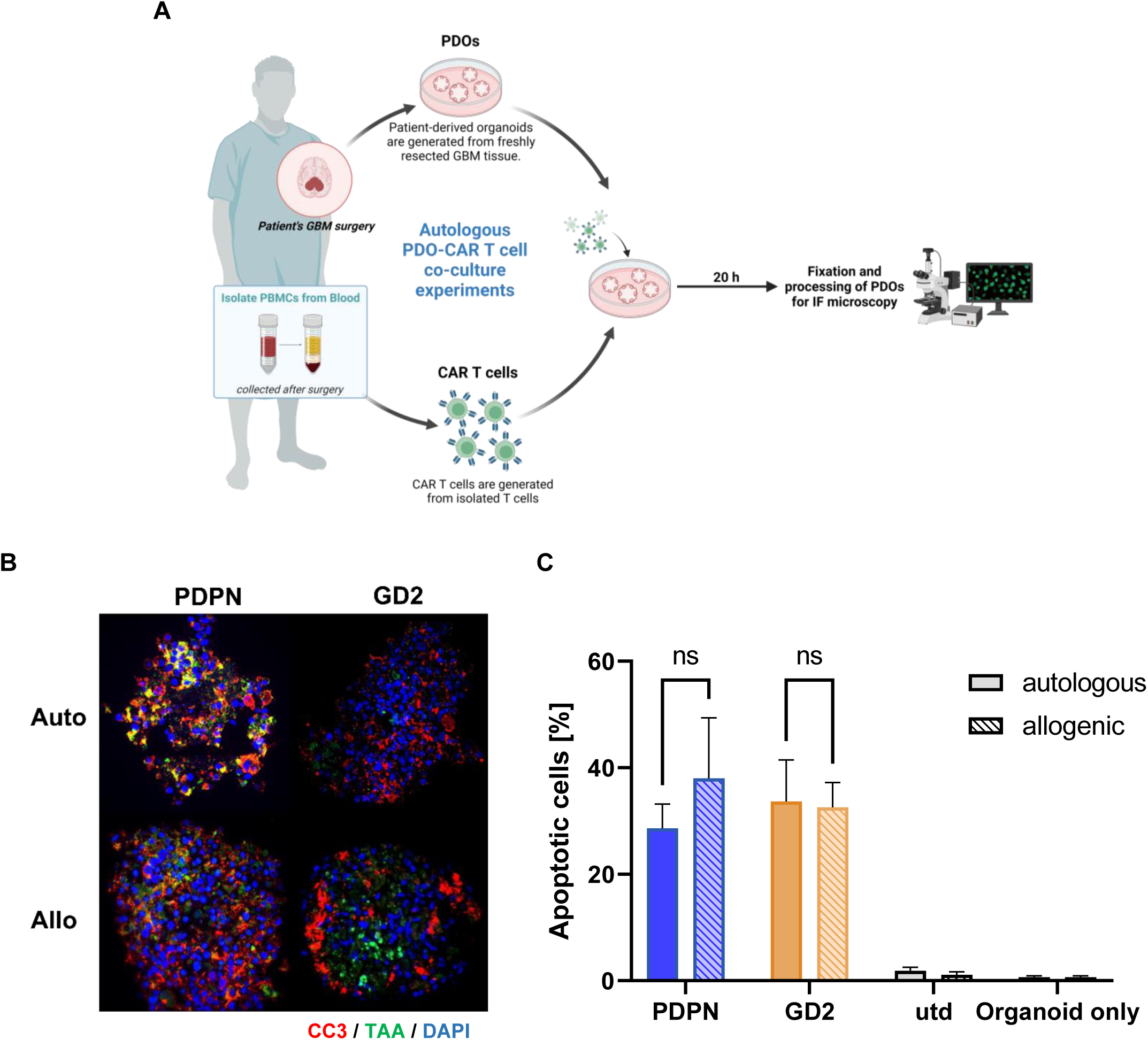
Autologous PDPN-CAR T cells are equally efficient in PDO co-cultures. **(A)** After the patient’s GBM surgery and organoid generation, PBMCs were isolated from peripheral blood prior to the patient’s follow-up therapy. CAR T cells were generated and co-cultures were performed for 20 hours. Created with BioRender.com. **(B)** Representative IF images of tumor cell apoptosis (DAPI^+^/PDPN^+^/CC3^+^). **(C, D)** Quantification of apoptotic cells and IFNγ levels after incubation with allogeneic and autologous PDPN-CAR T cells. All experiments were performed in triplicates. Data are shown as mean + SEM. Magnification: 40x. Scale bar: 50 µm.

**Supplementary Figure S4:**
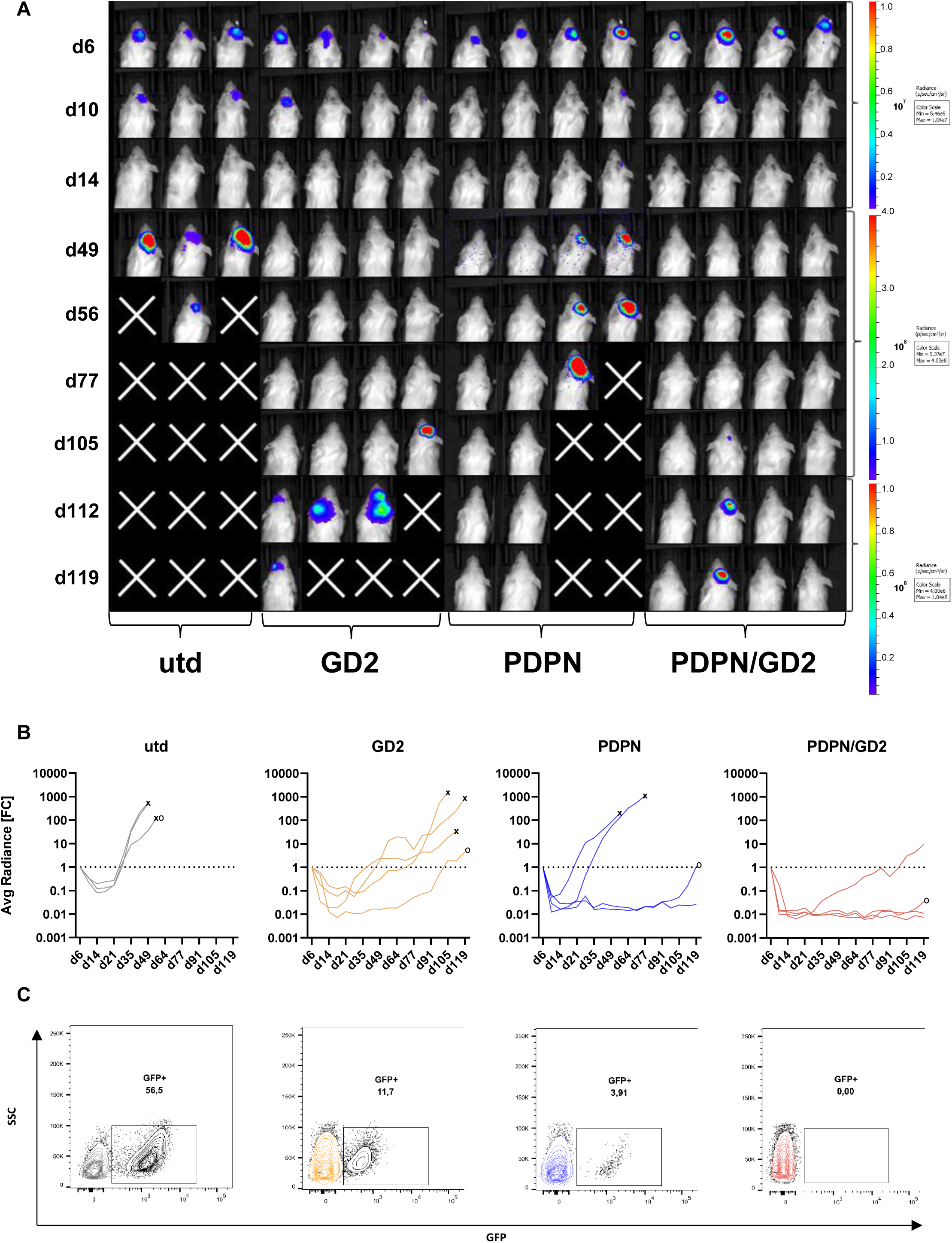
BLI images, fold change in average radiance and residual tumor cell analysis. **(A-B)** BLI imaging and fold change in average radiance. “x” indicates the time of euthanasia of mice according to the termination criteria; “o” indicates the mice shown below. **(C)** Representative flow cytometry plots of residual tumor cell analysis in the brain (GFP^+^ cells).

## REFERENCES

1. Thakkar JP, Dolecek TA, Horbinski C, et al. Epidemiologic and molecular prognostic review of glioblastoma. Cancer Epidemiol Biomarkers Prev. 2014;23(10):1985–1996. doi:10.1158/1055-9965.EPI-14-0275

2. Marton E, Giordan E, Siddi F, et al. Over ten years overall survival in glioblastoma: A different disease? J Neurol Sci. 2020;408:116518. doi:10.1016/J.JNS.2019.116518

3. Stupp R, Mason WP, van den Bent MJ, et al. Radiotherapy plus concomitant and adjuvant temozolomide for glioblastoma. N Engl J Med. 2005;352(10):987–996. doi:10.1056/NEJMOA043330

4. Gately L, McLachlan S, Dowling A, Philip J. Life beyond a diagnosis of glioblastoma: a systematic review of the literature. Journal of Cancer Survivorship. 2017;11(4):447–452. doi:10.1007/S11764-017-0602-7/TABLES/2

5. Omuro A, DeAngelis LM. Glioblastoma and Other Malignant Gliomas: A Clinical Review. JAMA. 2013;310(17):1842–1850. doi:10.1001/JAMA.2013.280319

6. Stupp R, Taillibert S, Kanner A, et al. Effect of Tumor-Treating Fields Plus Maintenance Temozolomide vs Maintenance Temozolomide Alone on Survival in Patients With Glioblastoma: A Randomized Clinical Trial. JAMA. 2017;318(23):2306–2316. doi:10.1001/JAMA.2017.18718

7. Cappell KM, Kochenderfer JN. Long-term outcomes following CAR T cell therapy: what we know so far. Nature Reviews Clinical Oncology 2023 20:6. 2023;20(6):359–371. doi:10.1038/s41571-023-00754-1

8. June CH, Sadelain M. Chimeric Antigen Receptor Therapy. N Engl J Med. 2018;379(1):64–73. doi:10.1056/NEJMRA1706169

9. Rodríguez-Lobato LG, Ganzetti M, Fernández de Larrea C, Hudecek M, Einsele H, Danhof S. CAR T-Cells in Multiple Myeloma: State of the Art and Future Directions. Front Oncol. 2020;10:541019. doi:10.3389/FONC.2020.01243/BIBTEX

10. Hou AJ, Chen LC, Chen YY. Navigating CAR-T cells through the solid-tumour microenvironment. Nat Rev Drug Discov. 2021;20(7):531–550. doi:10.1038/S41573-021-00189-2

11. Bagley SJ, O’Rourke DM. Clinical investigation of CAR T cells for solid tumors: Lessons learned and future directions. Pharmacol Ther. 2020;205. doi:10.1016/J.PHARMTHERA.2019.107419

12. O’Rourke DM, Nasrallah MP, Desai A, et al. A single dose of peripherally infused EGFRvIII-directed CAR T cells mediates antigen loss and induces adaptive resistance in patients with recurrent glioblastoma. Sci Transl Med. 2017;9(399). doi:10.1126/SCITRANSLMED.AAA0984

13. Ahmed N, Brawley V, Hegde M, et al. HER2-Specific Chimeric Antigen Receptor-Modified Virus-Specific T Cells for Progressive Glioblastoma: A Phase 1 Dose-Escalation Trial. JAMA Oncol. 2017;3(8):1094–1101. doi:10.1001/JAMAONCOL.2017.0184

14. Brown CE, Badie B, Barish ME, et al. Bioactivity and Safety of IL13Rα2-Redirected Chimeric Antigen Receptor CD8+ T Cells in Patients with Recurrent Glioblastoma. Clin Cancer Res. 2015;21(18):4062–4072. doi:10.1158/1078-0432.CCR-15-0428

15. Sasano T, Gonzalez-Delgado R, Muñoz NM, et al. Podoplanin promotes tumor growth, platelet aggregation, and venous thrombosis in murine models of ovarian cancer. J Thromb Haemost. 2022;20(1):104–114. doi:10.1111/JTH.15544

16. Astarita JL, Acton SE, Turley SJ. Podoplanin: emerging functions in development, the immune system, and cancer. Front Immunol. 2012;3(SEP). doi:10.3389/FIMMU.2012.00283

17. Quintanilla M, Montero LM, Renart J, Villar EM. Podoplanin in Inflammation and Cancer. Int J Mol Sci. 2019;20(3). doi:10.3390/IJMS20030707

18. Suzuki M, Cheung NK V. Disialoganglioside GD2 as a therapeutic target for human diseases. Expert Opin Ther Targets. 2015;19(3):349–362. doi:10.1517/14728222.2014.986459

19. Majzner RG, Ramakrishna S, Yeom KW, et al. GD2-CAR T cell therapy for H3K27M- mutated diffuse midline gliomas. Nature 2022 603:7903. 2022;603(7903):934–941. doi:10.1038/s41586-022-04489-4

20. Del Bufalo F, De Angelis B, Caruana I, et al. GD2-CART01 for Relapsed or Refractory High-Risk Neuroblastoma. New England Journal of Medicine. 2023;388(14):1284–1295. doi:10.1056/NEJMOA2210859/SUPPL_FILE/NEJMOA2210859_DATA-SHARING.PDF

21. Jacob F, Salinas RD, Zhang DY, et al. A Patient-Derived Glioblastoma Organoid Model and Biobank Recapitulates Inter- and Intra-tumoral Heterogeneity. Cell. 2020;180(1):188–204.e22. doi:10.1016/J.CELL.2019.11.036

22. Alsalkini M, Cibulková V, Breun M, et al. Cultivating Ex Vivo Patient-Derived Glioma Organoids Using a Tissue Chopper. J Vis Exp. 2024;2024(203). doi:10.3791/65952

23. Hudecek M, Schmitt TM, Baskar S, et al. The B-cell tumor–associated antigen ROR1 can be targeted with T cells modified to express a ROR1-specific chimeric antigen receptor. Blood. 2010;116(22):4532–4541. doi:10.1182/BLOOD-2010-05-283309

24. Schindelin J, Arganda-Carreras I, Frise E, et al. Fiji: an open-source platform for biological-image analysis. Nature Methods 2012 9:7. 2012;9(7):676–682. doi:10.1038/nmeth.2019

25. Love MI, Huber W, Anders S. Moderated estimation of fold change and dispersion for RNA-seq data with DESeq2. Genome Biol. 2014;15(12):1–21. doi:10.1186/S13059-014-0550-8/FIGURES/9

26. Sorokin M, Kholodenko I, Kalinovsky D, et al. RNA Sequencing-Based Identification of Ganglioside GD2-Positive Cancer Phenotype. Biomedicines. 2020;8(6). doi:10.3390/BIOMEDICINES8060142

27. Brown C, Alizadeh D, Starr R, … LW… EJ of, 2016 undefined. Regression of glioblastoma after chimeric antigen receptor T-cell therapy. Mass Medical SocCE Brown, D Alizadeh, R Starr, L Weng, JR Wagner, A Naranjo, JR Ostberg, MS BlanchardNew England Journal of Medicine, 2016•Mass Medical Soc. 2016;375(26):2561–2569. doi:10.1056/NEJMoa1610497

28. Durgin JS, Henderson F, Nasrallah MP, et al. Case Report: Prolonged Survival Following EGFRvIII CAR T Cell Treatment for Recurrent Glioblastoma. Front Oncol. 2021;11. doi:10.3389/FONC.2021.669071/FULL

29. Burger MC, Zhang C, Harter PN, et al. CAR-Engineered NK Cells for the Treatment of Glioblastoma: Turning Innate Effectors Into Precision Tools for Cancer Immunotherapy. Front Immunol. 2019;10. doi:10.3389/FIMMU.2019.02683/FULL

30. Mishima K, Kato Y, Kaneko MK, Nishikawa R, Hirose T, Matsutani M. Increased expression of podoplanin in malignant astrocytic tumors as a novel molecular marker of malignant progression. Acta Neuropathol. 2006;111(5):483–488. doi:10.1007/S00401-006-0063-Y

31. Martín-Villar E, Megías D, Castel S, Yurrita MM, Vilaró S, Quintanilla M. Podoplanin binds ERM proteins to activate RhoA and promote epithelial-mesenchymal transition. J Cell Sci. 2006;119(Pt 21):4541–4553. doi:10.1242/JCS.03218

32. Fehon RG, McClatchey AI, Bretscher A. Organizing the cell cortex: the role of ERM proteins. Nature Reviews Molecular Cell Biology 2010 11:4. 2010;11(4):276–287. doi:10.1038/nrm2866

33. Modrek AS, Eskilsson E, Ezhilarasan R, et al. PDPN marks a subset of aggressive and radiation-resistant glioblastoma cells. Front Oncol. 2022;12:941657. doi:10.3389/FONC.2022.941657/BIBTEX

34. Wu M, Shi Y, Liu Y, et al. Exosome-transmitted podoplanin promotes tumor-associated macrophage-mediated immune tolerance in glioblastoma. CNS Neurosci Ther. 2024;30(3). doi:10.1111/CNS.14643

35. Pula B, Witkiewicz W, Dziegiel P, Podhorska-Okolow M. Significance of podoplanin expression in cancer-associated fibroblasts: A comprehensive review. Int J Oncol. 2013;42(6):1849–1857. doi:10.3892/IJO.2013.1887/HTML

36. Shiina S, Ohno M, Ohka F, et al. CAR T Cells Targeting Podoplanin Reduce Orthotopic Glioblastomas in Mouse Brains. doi:10.1158/2326-6066.CIR-15-0060

37. Chalise L, Kato A, Ohno M, et al. Efficacy of cancer-specific anti-podoplanin CAR-T cells and oncolytic herpes virus G47Δ combination therapy against glioblastoma. Mol Ther Oncolytics. 2022;26:265–274. doi:10.1016/J.OMTO.2022.07.006

38. Kailayangiri S, Altvater B, Meltzer J, et al. The ganglioside antigen G(D2) is surface-expressed in Ewing sarcoma and allows for MHC-independent immune targeting. Br J Cancer. 2012;106(6):1123–1133. doi:10.1038/BJC.2012.57

39. Lammie GA, Cheung NKV, Gerald W, Rosenblum M, Cordon-Cardo C. Ganglioside gd(2) expression in the human nervous-system and in neuroblastomas - an immunohistochemical study. Int J Oncol. 1993;3(5):909–915. doi:10.3892/IJO.3.5.909

40. Chang HR, Cordon-Cardo C, Houghton AN, Cheung NK V, Brennan MF. Expression of Disialogangliosides GO2 and GD3 on Human Soft Tissue Sarcomas. doi:10.1002/1097-0142

41. Mennel HD, Bosslet K, Wiegandt H, Sedlacek HH, Bauer BL, Rodden AF. Expression of GD2-epitopes in human intracranial tumors and normal brain. Exp Toxicol Pathol. 1992;44(6):317–324. doi:10.1016/S0940-2993(11)80218-6

42. Furukawa K, Hamamura K, Ohkawa Y, Ohmi Y, Furukawa K. Disialyl gangliosides enhance tumor phenotypes with differential modalities. Glycoconj J. 2012;29(8-9):579–584. doi:10.1007/S10719-012-9423-0

43. Majzner RG, Ramakrishna S, Yeom KW, et al. GD2-CAR T cell therapy for H3K27M- mutated diffuse midline gliomas. Nature 2022 603:7903. 2022;603(7903):934–941. doi:10.1038/s41586-022-04489-4

44. Wang N, Hu X, Cao W, et al. Efficacy and safety of CAR19/22 T-cell cocktail therapy in patients with refractory/relapsed B-cell malignancies. Blood. 2020;135(1):17–27. doi:10.1182/BLOOD.2019000017

45. Schultz LM, Muffly LS, Spiegel JY, et al. Phase I Trial Using CD19/CD22 Bispecific CAR T Cells in Pediatric and Adult Acute Lymphoblastic Leukemia (ALL). Blood. 2019;134(Supplement_1):744–744. doi:10.1182/BLOOD-2019-129411

46. Tousley AM, Rotiroti MC, Labanieh L, et al. Co-opting signalling molecules enables logic-gated control of CAR T cells. Nature 2023 615:7952. 2023;615(7952):507–516. doi:10.1038/s41586-023-05778-2

47. Brown CE, Alizadeh D, Starr R, et al. Regression of Glioblastoma after Chimeric Antigen Receptor T-Cell Therapy. N Engl J Med. 2016;375(26):2561–2569. doi:10.1056/NEJMOA1610497

48. Choi BD, Suryadevara CM, Gedeon PC, et al. Intracerebral delivery of a third generation EGFRvIII-specific chimeric antigen receptor is efficacious against human glioma. J Clin Neurosci. 2014;21(1):189–190. doi:10.1016/J.JOCN.2013.03.012

49. Brown CE, Hibbard JC, Alizadeh D, et al. Locoregional delivery of IL-13Rα2-targeting CAR-T cells in recurrent high-grade glioma: a phase 1 trial. Nature Medicine 2024 30:4. 2024;30(4):1001–1012. doi:10.1038/s41591-024-02875-1

50. Prapa M, Chiavelli C, Golinelli G, et al. GD2 CAR T cells against human glioblastoma. npj Precision Oncology 2021 5:1. 2021;5(1):1–14. doi:10.1038/s41698-021-00233-9

